# Deep-Learning Cortical Registration Guided by Structural and Diffusion MRI and Connectivity

**DOI:** 10.1101/2025.10.31.685908

**Authors:** Zhen Zhou, Jian Li, Jonathan Williams, Bruce Fischl, Iman Aganj

**Affiliations:** Athinoula A. Martinos Center for Biomedical Imaging, Massachusetts General Hospital, Harvard Medical School, Boston, Massachusetts, USA

**Keywords:** Cortical surface registration, heat diffusion smoothing, semi-supervised learning, functional alignment

## Abstract

Accurate cortical surface registration is crucial for group-level neuroimaging analyses, yet geometry-based methods often yield suboptimal functional alignment across subjects due to substantial inter-individual variability. We present a novel deep-learning approach that incorporates white matter structural connectivity features from diffusion MRI (dMRI) tractography into the Joint Surface-based Registration and Atlas Construction (JOSA) framework. Our method generates vertex-wise connectivity maps by detecting streamline-surface intersections, followed by heat diffusion smoothing on the cortical manifold. We combine these connectivity features with scalar diffusion metrics (fractional anisotropy and apparent diffusion coefficient) and structural features as input to JOSA (which we call “JOSAConn”). Evaluated on HCP-YA subjects across 15 task contrasts, our method significantly outperforms FreeSurfer in functional alignment (Bonferroni-corrected *p* < 0.05 for 12 of 15 contrasts). This multimodal approach demonstrates that structural connectivity effectively bridges the gap between cortical geometry and functional organization while maintaining clinical applicability.

## 1. INTRODUCTION

Accurate cortical surface registration is essential for enabling group-level analyses in neuroimaging studies. Traditional surface-based registration methods like FreeSurfer [1] rely primarily on geometric features such as sulcal depth and cortical curvature. Recent advances in connectivity-based parcellation [2-5] have demonstrated that structural and functional connectivity patterns – computed via diffusion MRI (dMRI) and functional MRI (fMRI) – provide complementary information about cortical organization that is not fully captured by geometry alone. Structural connectivity has also been leveraged for cortical registration [6-8].

We propose a novel approach that incorporates dMRI-derived structural connectivity features into cortical surface registration. Specifically, we leverage the Joint Surface-based Registration and Atlas Construction (JOSA) [9, 10] framework in a semi-supervised learning paradigm. JOSA jointly learns to register cortical surfaces and construct population atlases by optimizing registration using both anatomical features and task-based functional activation patterns. In this work, we extend JOSA by incorporating white matter connectivity features derived from dMRI tractography as additional anatomical constraints, which we will call “JOSAConn”. This approach aims to improve functional alignment by ensuring that regions with similar structural connectivity profiles are registered together, even when their geometric features differ.

In this work, we generate vertex-wise structural connectivity features by identifying where tractography streamlines intersect the cortical surface and then applying heat diffusion smoothing on the surface to produce spatially coherent connectivity maps. The connectivity and traditional anatomical features, along with functional task data in a semi-supervised learning framework, guide the registration to align both structural connectivity and functional organization across subjects. Our results demonstrate our ability to achieve a superior functional registration performance compared to FreeSurfer using the proposed approach, but without requiring fMRI data at inference.

In the following, we describe the proposed method in detail (Section 2) and present experimental results (Section 3) along with some concluding remarks (Section 4).

## 2. METHODS

### 2.1. Data and Processing

We used structural MRI (sMRI), dMRI, and fMRI data of 1,064 subjects from the Human Connectome Project - Young Adult (HCP-YA) dataset [11]. We processed the sMRI data using FreeSurfer v7.4.0 and reconstructed the subjects’ surfaces from the T1-weighted images (default `recon-all` parameters).

We then ran the FreeSurfer dMRI processing pipeline, which also incorporates commands from FSL [12]. This process involved propagating 85 automatically segmented cortical and subcortical regions from the sMRI to the dMRI space using boundary-based image registration [13]. Next, we used our publicly available dMRI analysis toolbox (www.nitrc.org/projects/csaodf-hough) to reconstruct the diffusion orientation distribution function in constant solid angle (CSA-ODF) [14] from the b=3000 s/mm2 q-shell and run Hough-transform global probabilistic tractography [15] to generate an optimal (highest-score) streamline passing through each of the 10,000 seed points for each subject. Fractional anisotropy (FA) and apparent diffusion coefficient (ADC) values were also extracted from the diffusion tensor imaging model fitted to the multi-shell diffusion data.

We processed HCP’s task fMRI data using FreeSurfer’s FsFast pipeline for functional analyses. We applied a 2 mm FWHM surface-based Gaussian smoothing to avoid blurring signals across sulci/gyri. Task fMRI data from 7 HCP tasks (emotion processing, gambling, language, motor, social cognition, relational processing, and working memory) were used. Task contrast maps (*t*-statistics) were computed for each subject through general linear modeling using their corresponding block designs. We selected 15 unique and representative contrasts (faces-shapes, punish-reward, story-math, lf-avg, lh-avg, rf-avg, rh-avg, t-avg, match-rel, random-tom, 2bk-0bk, body-avg, face-avg, place-avg, and tool-avg) from the 7 tasks for evaluation purposes [16]. These functional activation patterns provided ground truth for training the registration network to align regions with similar functional profiles, while the structural and diffusion features enabled the network to generalize to new subjects without requiring task fMRI at inference time.

### 2.2. Structural and Connectivity Feature Generation

We extracted surface-based structural connectivity features from dMRI and sMRI data of the HCP-YA [11] subjects. The analysis generated vertex-wise connectivity maps on cortical surfaces by integrating white-matter tractography with FreeSurfer-reconstructed anatomical surfaces. The resulting features capture the spatial distribution of white matter connections terminating at each cortical location, providing a high-resolution representation of brain structural connectivity suitable for subsequent statistical analysis.

We identified where tractography streamlines intersect the cortical (white/gray) surface using a vectorized algorithm based on the Möller-Trumbore ray-triangle intersection test. Each streamline was decomposed into consecutive line segments, and we computed intersections between all segment-triangle pairs accelerated on the graphics processing unit. For each potential interaction, we validated intersections based on geometric constraints to ensure non-parallel configurations and that interactions occurred within segment bounds. We transformed the discrete intersection data into a continuous vertex-wise connectivity feature vector through a multi-step aggregation process. The feature vector had 85 elements, each corresponding to a segmented (cortical or subcortical) region of interest (ROI). First, streamline scores were accumulated at the mesh *face* level, with each element of the feature vector receiving scores from streamlines passing through its corresponding ROI. Face-level scores were then projected to vertices by distributing each face’s aggregated score equally to its three constituent vertices, ensuring connectivity information from all intersecting streamlines contributed to nearby cortical locations. To reduce the right-skewed distribution typical of connectivity data [17], we transformed the vertex scores as *s*← log(1 + *s*), producing features more amenable to parametric statistical analyses.

We then spatially smoothed the features on the triangular mesh manifold by applying the heat equation to the feature maps. We solved the heat equation on the surface using an implicit time-stepping scheme, which naturally accounts for local surface geometry and provides numerically stable smoothing. Following the heat diffusion framework, we model the smoothing of vertex connectivity scores as a heat flow process on the cortical surface manifold. The heat equation describes how temperature (in our case, connectivity scores) diffuses over time based on local gradients:

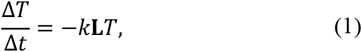

where *T*is the vertex-wise temperature (here, the connectivity score, *s*) vector, Δ*t* is the time step, *k*is the diffusion coefficient (*k*=1 in our implementation), **L** =**D**− **A** is the graph Laplacian matrix, with **D**and **A** the graph’s degree and adjacency matrices, respectively. We used simple forward Euler time-stepping [18] with Δ*t* =0.001 for *n*_*max*_ =100 iterations to solve the heat equation.

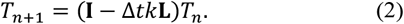

This procedure yielded the final vertex-wise connectivity features for each hemisphere, representing spatially smoothed estimates of white matter connectivity density at approximately 150,000 cortical locations per subject. The final output for each subject consisted of two feature matrices (left and right hemispheres), each with dimensions ∼150,000 × 85. Rows represent cortical vertices and columns represent anatomical ROIs, with each element encoding the smoothed structural connectivity strength for a specific vertex-ROI pair.

Using FreeSurfer’s `mri_vol2surf` command, we sampled FA and ADC values (from dMRI images) and T1- and T2-weighted intensity values (from sMRI images) at each cortical vertex by projecting them to mid-cortical depth. These scalar features provide complementary information about local tissue properties: FA reflects fiber integrity, ADC indicates water diffusivity, and T1 and T2 intensities capture gray matter contrast characteristics. The resulting vertex-wise scalar maps were combined with structural connectivity features and geometric features (sulcal depth, curvature, cortical thickness) derived from the FreeSurfer pipeline [1] as multimodal input to the JOSAConn registration framework.

### 2.3. Semi-supervised Registration using JOSAConn

We leveraged the JOSA framework [9, 10] for accurate cortical registration. As the main input to JOSAConn, we used sMRI features (sulcal depth, curvature, thickness, T1 intensity, T2 intensity), dMRI scalar features (FA, ADC), and dMRI connectivity features (as described in Section 2.2). We used the *t* maps derived from task fMRI in the semi-supervised path in JOSAConn. This semi-supervised training strategy is of critical importance because it allows the network to learn local structural/diffusion features that are predictive of the brain function, but without the need for the task fMRI data to be available at inference. All features are rigidly rotated to the `fsaverage` space and parameterized into 2D flat maps.

We implemented JOSA based on VoxelMorph [19, 20] with 5 encoding and decoding layers in the U-Net with a flat 128 channels. We used an MSE loss for the data fidelity, and a gradient loss on both the joint and the modality-specific deformation fields to encourage smooth warps. We also applied a Jacobian loss on the composited deformation fields to avoid potential topological errors during the warping. To account for the difference in sampling density during parameterization, we performed prior and distortion corrections for all losses [10, 21]. To account for the discontinuity across the boundaries in the parameterized images, we applied a spherical padding with a circular padding on the left and right, and a reflection after a *π*shift on the top and bottom of the image. We employed a two-stage learning procedure via learning the atlases first and then learning the deformation fields with fixed atlases. We also evaluated the losses bidirectionally, both in the atlas and the subject space, to help avoid atlas expansion or drift [22]. We used a batch size of 8, augmenting the data for each batch of input using both random warps and random noise. We employed the Adam optimizer with an initial learning rate of 10^-3^ for the first 500 epochs, after which it decreased by a factor of 0.98 if the validation loss did not decrease after every 100 epochs. The regularization weights between the joint warp and the separate warps were empirically set to 0.1 and 0.2, respectively (relative to the weight of 1 for the data fidelity term). We used TensorFlow [23], Keras [24], and Neurite [25] for implementations. We held the 100 unrelated subjects (provided by HCP-YA) for testing and randomly split the remaining subjects into a training set (800 subjects) and a validation set (164 subjects).

### 2.4. Comparison and Evaluation

We compared our JOSAConn with FreeSurfer. For both methods, we performed a qualitative evaluation by computing the group mean task maps over the 100 testing subjects who were unseen during the training. We note that these task maps were registered to the atlas space using the predicted task-dedicated deformation (the semi-supervised path), which is, however, solely determined by the subject’s geometric and diffusion data. We mapped the mean images back to the surface space, overlaying with the curvature map, and visualized them using Freeview.

Quantitatively, we then evaluated the registration performance using Pearson’s correlation between the registered individual task map and the group mean for each task contrast separately. We present the distribution of the correlation across 100 testing subjects, testing the difference between JOSAConn and FreeSurfer using a Wilcoxon signed rank test, as these are paired measures.

## 3. RESULTS AND DISCUSSION

### 3.1. Generated Structural and Diffusion Features

In **Fig. 1**, the right hemisphere cortical surface displays vertex-wise connectivity scores derived from the intersections of white matter streamlines with the FreeSurfer-reconstructed surface. Vertex-wise connectivity features were generated through our processing pipeline: streamline-surface intersection detection, face-level score aggregation for brainstem-connected pathways, vertex projection via neighborhood averaging, and 100 iterations of heat diffusion smoothing on the triangular mesh manifold. This multi-view visualization demonstrates the spatial structure of brainstem connectivity feature that, combined with connectivity features from other regions, as well as geometric and scalar diffusion metrics, serve as multimodal input to the registration network. The connectivity distribution shows expected anatomical organization with peak intensities (red-yellow) in primary motor and sensory cortices, consistent with corticospinal, corticobulbar, and ascending sensory tract anatomy.

**Fig. 1.**
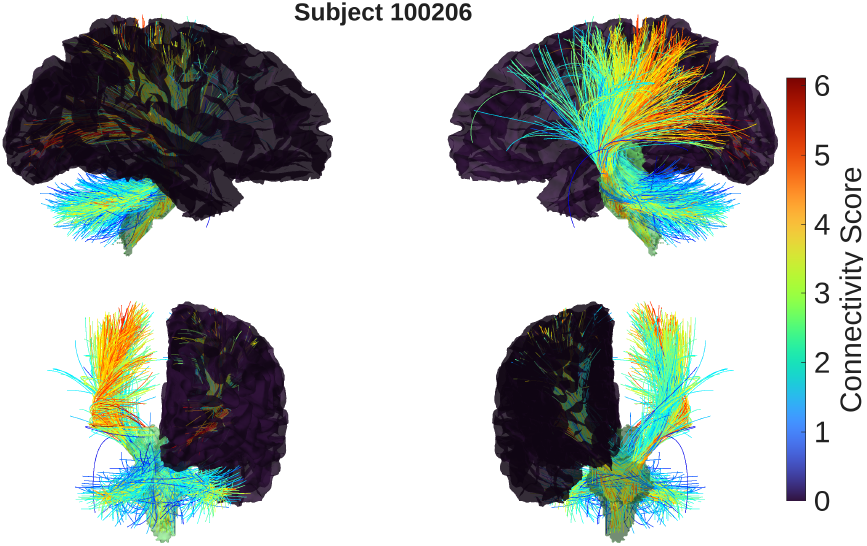
Brainstem structural connectivity and streamline visualization. This figure illustrates all tractography streamlines going through the brainstem of an HCP-YA subject, visualized from four views. The intersections of streamlines with cortical surfaces are color-coded by connectivity scores, overlaid on FreeSurfer-reconstructed cortical surfaces.

### 3.2. JOSAConn Registration Results

**Fig. 3** qualitatively compares the task registration performance using the HCP language task story-math contrast as an example. The group mean of the FreeSurfer-registered language *t* maps overlaid on top of the corresponding group mean curvature map is shown on the left, with the JOSAConn counterpart shown on the right. JOSAConn exhibits a stronger language-area activation and sharper transition from language-active regions to language-inactive regions in comparison to FreeSurfer, particularly in regions in BA 44/45 (Broca’s area) and the frontal eye field.

**Fig. 2** shows a quantitative comparison of the registration performance using correlations between individual task *t* maps and their corresponding group means. (The higher the correlation, the better the registration.) Across 15 unique contrasts from 7 HCP tasks, JOSAConn demonstrated superior functional alignment in 14 contrasts compared to FreeSurfer. Wilcoxon signed-rank tests revealed that 12 of these 14 contrasts achieved statistical significance with (Bonferroni-corrected) *p* < 0.05, while 2 contrasts showed improvement without reaching significance. Only the rf-avg (right foot vs. average) motor contrast showed slightly lower performance for JOSAConn compared to FreeSurfer, though this difference was not statistically significant.

**Fig. 2.**
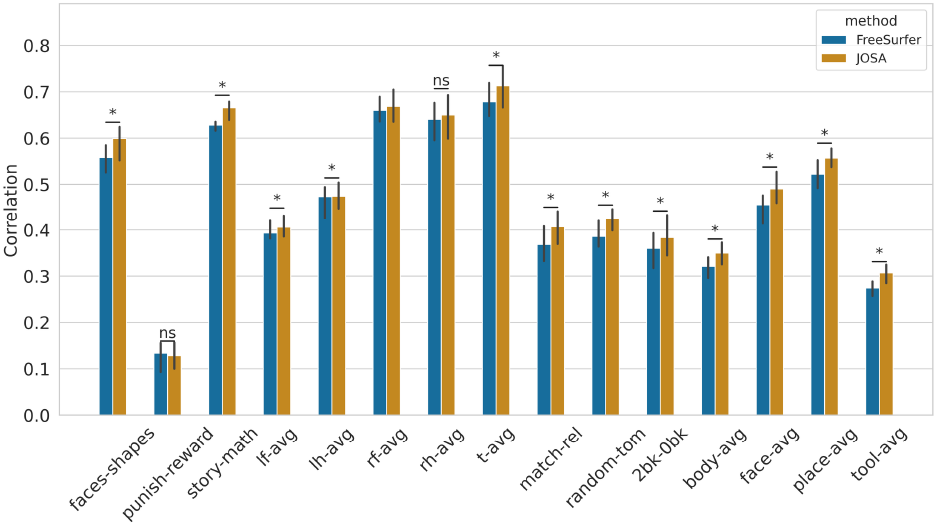
Quantitative comparison of registration performance. The correlations between the individual *t* maps of the 100 testing subjects and their corresponding group mean are bar-plotted. The *x*-axis lists the selected 15 unique contrasts across 7 HCP tasks. FreeSurfer results are shown in blue, and the results of the JOSAConn are shown in orange. Each bar represents the median correlation across 100 test subjects, with vertical black lines indicating 95% confidence intervals. JOSAConn outperforms FreeSurfer in 14 of 15 contrasts, with 12 showing statistically significant improvement (marked with ‘*’, Wilcoxon signed-rank test, *p* < 0.05) and 2 showing non-significant improvement (marked with ‘ns’, *p* > 0.05).

**Fig. 3.**
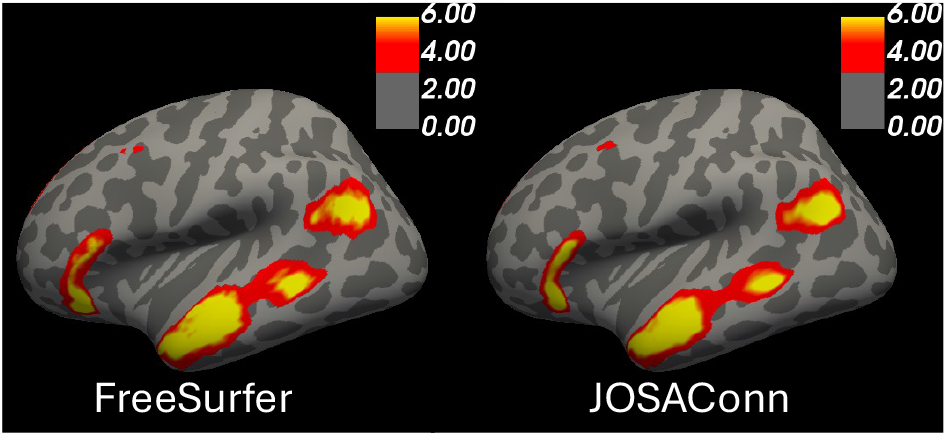
Qualitative comparison of registration performance. The group mean curvature map over the 100 testing subjects is shown in gray for FreeSurfer (left) and JOSAConn (right). The corresponding group mean *t*-maps for the language task story-math contrast are overlaid on top with heat colormap.

## 4. CONCLUSIONS

We have introduced a new approach to cortical surface registration that incorporates dMRI-derived structural connectivity features into the JOSA framework using semi-supervised learning. By combining tractography-based connectivity maps with dMRI scalar diffusion features (FA and ADC) as well as sMRI features (sulcal depth, curvature, thickness, T1 intensity, T2 intensity) derived from the standard FreeSurfer pipeline, our method achieved significantly improved functional alignment compared to traditional geometry-based registration. In our quantitative evaluation on 15 task contrasts across 7 HCP tasks, our JOSAConn significantly outperformed FreeSurfer registration (for 12 of 15 contrasts). Importantly, the semi-supervised training strategy enables inference using only structural and diffusion data, without requiring task fMRI. This multimodal approach provides a principled framework for improving cortical registration by leveraging complementary information from white matter connectivity patterns.

## COMPLIANCE WITH ETHICAL STANDARDS

The methodology presented in this article was evaluated retrospectively on publicly available de-identified images from the HCP-YA database [11], thereby not requiring ethical approval.

## ACKNOWLEDGMENTS

Support for this research was provided by the National Institutes of Health (NIH) grants R01AG068261, RF1NS128961, R01NS130119, and R01EB023281, and the Michael J. Fox Foundation for Parkinson’s Research (MJFF-021226). Computational resources were provided through the Massachusetts Life Sciences Center.

Additional support was provided by the BRAIN Initiative Cell Atlas Network (BICAN) grants U01MH117023, UM1MH134812 and UM1MH130981, the Brain Initiative Brain Connects consortium (U01NS132181, 1UM1NS132358-01), the National Institute for Biomedical Imaging and Bioengineering (R21EB018907, R01EB019956, P41EB030006), the National Institute on Aging (R21AG082082, 1R01AG064027, R01AG016495, 1R01AG070988), the National Institute of Mental Health (UM1MH130981, R01MH123195, R01MH121885, 1RF1MH123195), the National Institute for Neurological Disorders and Stroke, (1U24NS135561-01, R01NS070963, 2R01NS083534, R01NS105820, R25NS125599), and was made possible by the resources provided by Shared Instrumentation Grants 1S10RR023401, 1S10RR019307 and 1S10RR023043. Additional support was provided by the NIH Blueprint for Neuroscience Research (5U01-MH093765), part of the multi-institutional Human Connectome Project.

B. Fischl is a medical advisor to DeepHealth, a company whose medical pursuits focus on brain imaging and measurement technologies. His interests were reviewed and are managed by Massachusetts General Hospital and Mass General Brigham in accordance with their conflict-of-interest policies.

## Notes

http://www.nitrc.org/projects/csaodf-hough

https://silencer1127.github.io/software/JOSA/josa_main

### Summary of Updates

Update the results with new codes.

